# Base-mismatched crRNA in CRISPR Enabled Universal One-pot Rapid Nucleic Acid Detection

**DOI:** 10.64898/2026.09.30.755528

**Authors:** Xu Jianhao, Wang Xin, Yuan Bing, Hu Mingda, Xin Wenwen, Wang Jinglin, Ren Hongguang, Yuan Yuan

## Abstract

CRISPR-Cas12a-based diagnostics are promising for field detection yet are often constrained by two-step workflows or laborious optimization of one-pot reactions. Herein, we present a universal one-pot RPA/CRISPR-Cas12a strategy using base-mismatched crRNA in the seed region (positions 1–6). Introducing of single-nucleotide mismatches (complementary or error) into crRNA suppresses cis-cleavage activity, enabling amplification and detection in a single tube without PAM re-design or costly modified crRNA. Molecular dynamics (MD) simulations indicated that moderate potential energy (PE) variation correlates with improved screening performance, providing guidance for the selection of effective mismatches. This strategy successfully converted established two-step assays for *Burkholderia pseudomallei* and *Brucella* into one-pot formats, achieving detection of targets at 10 copies/μL within 30 min with good specificity. Lyophilized pellets supported field testing on lithium-ion battery-powered devices, and experiments with simulated soil samples confirm the robustness of the approach. Together, this cost-effective, adaptable strategy can streamline RPA-CRISPR diagnostics for infectious disease surveillance.

## Introduction

CRISPR-Cas12a-based diagnostic platforms are considered promising candidate technologies for field or home-based detection of infectious diseases due to their isothermal, rapid, and highly adaptable characteristics^1, 2^. Classical CRISPR diagnostic methods employ a two-step approach, in which nucleic acid amplification (NAA) and CRISPR detection reactions are performed sequentially. These methods offer advantages such as high sensitivity and excellent specificity and have been extensively reported in the literature^3-5^. However, the two-step reaction not only complicates the operational procedure and increases the risk of aerosol contamination but also hinders integration with downstream detection devices, making it difficult to meet the portability requirements for field deployment^1, 6^.

Integrating NAA and CRISPR detection into a one-pot in vitro diagnostic format represents an effective strategy to overcome these limitations. However, suppressing cis-cleavage activity of CRISPR within the one-pot reaction—thereby enabling better accumulation of amplification products—remains a key challenge for achieving high-performance pathogen detection^6, 7^. Previous studies have shown that reducing the target binding affinity of crRNA by employing suboptimal PAMs, or using fully PAM-free target sequences for certain Cas-tolerant variants, can effectively suppress cis-cleavage and improve the sensitivity of one-pot assays^8, 9^. Nevertheless, for the widely reported PAM-dependent two-step assays, such design strategies require redesign of target sequences and crRNAs, incurring considerable time costs. Moreover, for strictly PAM-dependent Cas proteins such as Cas12b, the complete absence of a cognate PAM abolishes target binding and all cleavage activities, rendering PAM-free strategies entirely non-functional rather than just suppressive. This severely limits the generalizability of such methods^10^. Alternatively, other researchers have suggested using photo-modified crRNA to substantially abolish cis-cleavage, offering a universal one-pot detection strategy^11-13^. However, although structural optimization has been performed to reduce synthesis complexity, the cost of modified crRNA remains substantially higher than that of unmodified crRNA, thereby increasing the overall detection cost.

In addition to design strategies targeting PAM sequences or crRNA repeat regions, engineering the crRNA spacer sequence has also attracted attention and has been frequently applied in gene editing and SNP detection^14, 15^. In vivo investigations revealed that base mutations in the spacer seed region, which create mismatches against target DNA, substantially lower Cas12a cis-cleavage activity.^16^. Therefore, engineering of the crRNA seed region also holds potential for developing one-pot RPA/CRISPR-Cas12a in vitro diagnostic methods. However, the binding affinity between crRNA and target DNA varies with target sequences. To develop a more broadly applicable one-pot in vitro diagnostic assay, systematic data comparison and rational design guidance are still required.

Molecular dynamics (MD) computes atomistic trajectories by solving Newton’s equations of motion. Currently, MD simulations are used to investigate the molecular interaction mechanisms within CRISPR systems^17^. Use of MD simulations to characterize molecular interaction features may help elucidate how the type and position of base mutations within the crRNA seed region influence the trans-cleavage activity of Cas12a.

Based on our previously established two-step RPA/CRISPR-Cas12a methods for three critical biothreats (*Bacillus anthraci*s, *Brucella*, and *Burkholderia pseudomallei*), this study performed targeted base mutagenesis within the seed region (nucleotide positions 1–6) of crRNA (Figure 1A). By combining MD simulations with wet-lab experiments, the optimal types and positions of base mismatches were identified. Accordingly, we propose a crRNA base-mismatch strategy to construct a universal one-pot RPA/CRISPR-Cas12a diagnostic approach for the above targets (Figure 1B). This facile strategy enables the low-cost and efficient conversion of conventional two-step diagnostic workflows into simplified one-pot formats.

**Figure 1.**
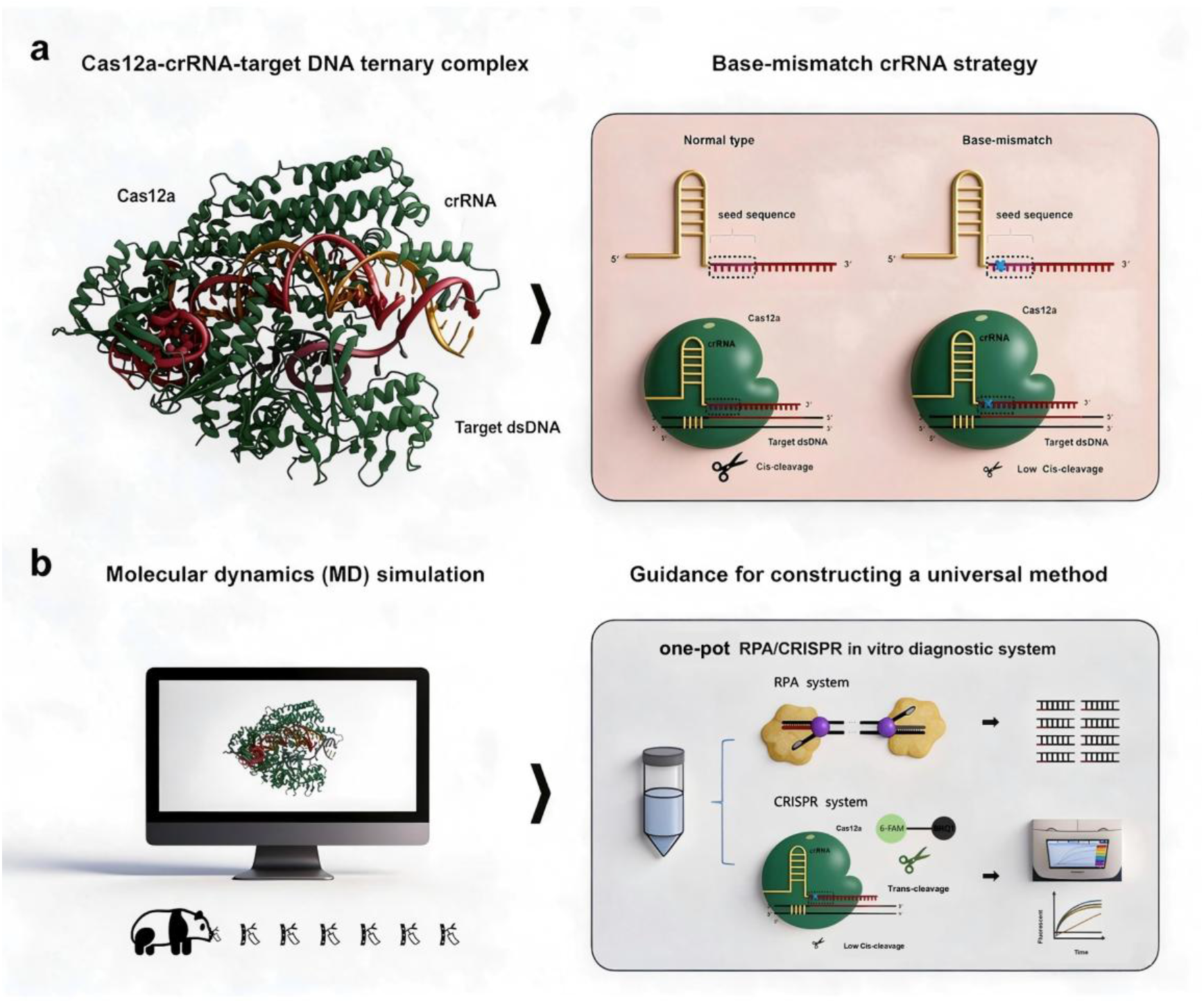
Schematic illustration of the study design. **(A)** Proposal and underlying principle of the base-mismatched crRNA strategy. **(B)** Application of molecular dynamics simulations to guide the establishment of a universal one-pot RPA/CRISPR-based in vitro diagnostic method.

## Methods

### Materials

heat-inactivated *Bacillus anthracis, Yersinia pestis, Francisella tularensis, Brucella, Burkholderia pseudomallei, Burkholderia mallei, Salmonella typhi, Vibrio cholerae, Staphylococcus aureus, Bacillus cereus, Bacillus subtilis, Bacillus thuringiensis* and their genomic DNA were provided by the State Key Laboratory of Pathogen and Biosecurity (Beijing, China).

The TwistAmp™ Basic Kit was purchased from TwistDX (Maidenhead, Berkshire, UK). The LbCas12a and RNase inhibitor were purchased from New England Biolabs Co (Ipswich, MA, USA). DNase/RNase-Free distilled, deionized water (ddH2O) provided by TIANGEN Biochemical Co., Ltd (Beijing, China). All the nucleic acid oligonucleotides used in this study were purchased from GenScript Biotech Corporation. (Nanjing, China), the RPA primers and crRNA were adopted from our previous studies, and the corresponding mismatch variants were designed accordingly. The sequences are shown in Table S1.

### Establishment of a one-pot RPA/CRISPR-Cas12a system

A one-pot RPA/CRISPR system for *B. anthracis* was established using the primers (*5345*-F, *5345*-R) and crRNA (*5345*-crRNA) from the two-step RPA/CRISPR method^18^. The composition of the one-pot system was based on the study by Ma et al^19^. Subsequently, the conditions for crRNA concentration, Cas protein concentration, probe concentration, and reaction temperature were optimized sequentially. The specific components are listed in Table S2.

Primers (*LC1*-F, *LC1*-R) and crRNA (*LC1*-crRNA) from the two-step RPA/CRISPR detection method for *B. pseudomallei* ^20^, as well as the primers (*omp2a*-F, *omp2a*-R) and crRNA (*omp2a*-crRNA) from the two-step RPA/CRISPR detection method for *Brucella*^21^, were subsequently used with the same component concentrations as those established for the one-pot RPA/CRISPR-Cas12a system for *B. anthracis* (Table S2). The applicability of the established system to these additional pathogenic targets was then evaluated.

### Design principles of base-mismatched crRNA

Mutations were introduced into bases within the seed region (positions 1–6) of the crRNA spacer sequence, and two mismatch patterns were designed. The first was a complementary mismatch (Cmis), in which the base at a given position was mutated to its complementary base, such as adenine ribonucleotide (A) to uracil ribonucleotide (U) or guanine ribonucleotide (G) to cytosine ribonucleotide (C). The second was an error mismatch (Emis), in which the base at a given position was mutated to an unrelated base, such as A to C or G to U. According to these rules, both mutation types were introduced into the seed-region bases of *LC1*-crRNA and *omp2a*-crRNA. For each crRNA, 12 mutated variants were designed, yielding a total of 24 base-mismatched crRNAs. The designed crRNAs are listed in Table S1

### Molecular dynamics simulation methods

A subset of the base-mismatched crRNAs was subjected to MD simulations, and interaction characteristics among Cas12a, crRNA, and target DNA were calculated under varying conditions. The original structure was derived from a high-resolution crystal structure of the Cas12a/crRNA-DNA ternary complex lacking PAM-distal double-stranded DNA (PDB ID: 5XUS) (Figures 2A and 3A). The web-based 3DNA platform (http://web.x3dna.org/) was used to introduce nucleic acid base mutations, resulting in a total of 15 Cas12a/crRNA/target DNA complexes for MD simulations.

**Figure 2.**
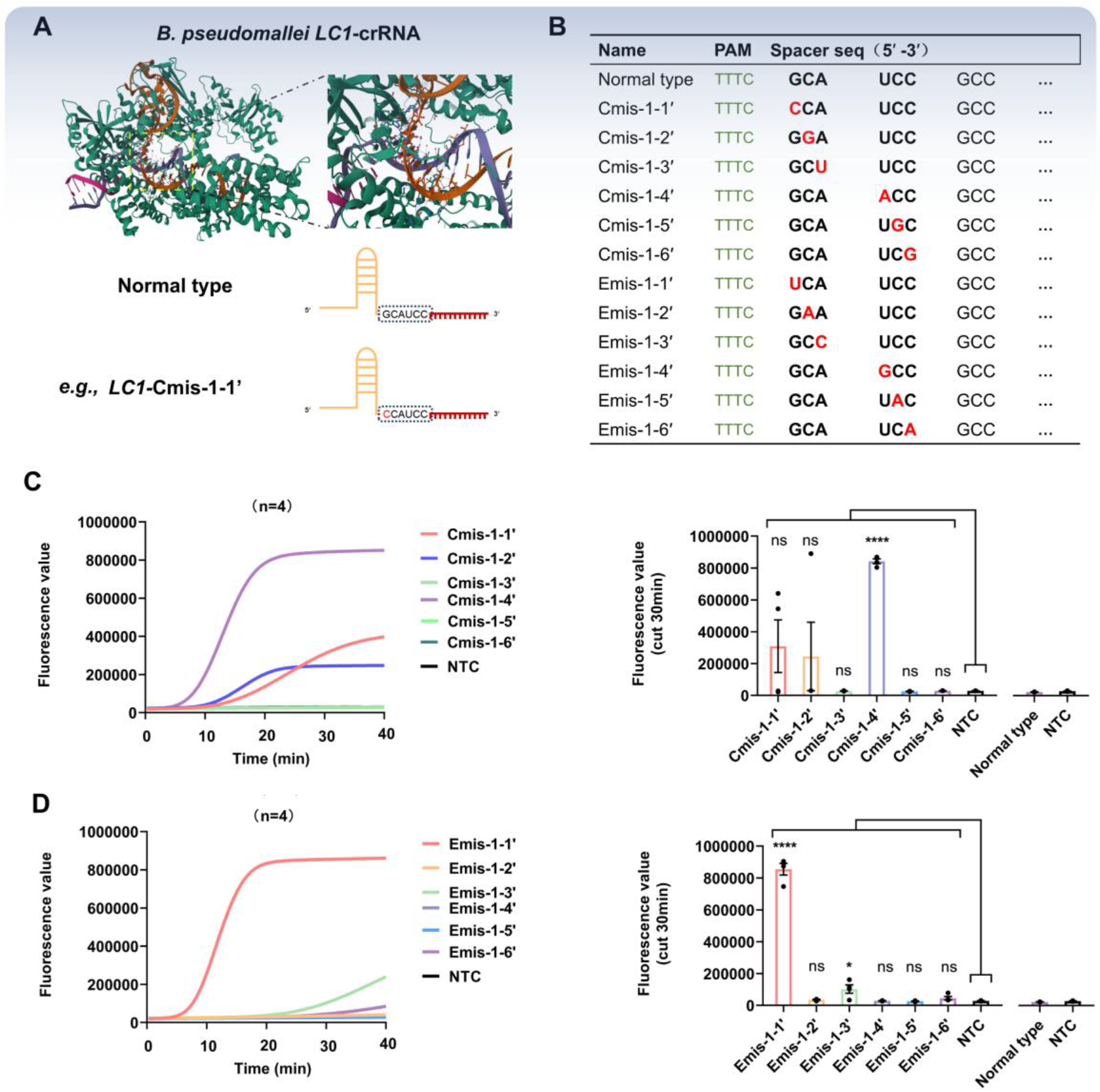
Screening results of base-mismatched crRNAs targeting *B. pseudomallei LC1*. Schematic illustration. **(B)** Mutation sites within the seed region of the 12 base-mismatched crRNAs. **(C)** Left panel: fluorescence amplification curves obtained from the Cmis validation group; right panel: bar chart showing the fluorescence signal values measured at the 30-min reaction time point. **(D)** Left panel: fluorescence amplification curves from the Emis validation group; right panel: bar chart of the fluorescence signal values at 30 min. (n = 4, mean ± SEM; statistical significance was assessed by *t*-test, with **** indicating *p* < 0.0001 and ns denoting not significant).

All MD simulations were performed using OpenMM (v8.0.0)^22^ with the Amber14 force field suite^23^. Specifically, amber14/protein.ff14SB, amber14/DNA.OL15, and amber14/RNA.OL3 were adopted for parametrization of proteins, DNA, and RNA, respectively, and the TIP3P water model (amber14/tip3pfb) was employed for solvent representation^24^. Simulations were carried out in 0.15 M KCl solution, a 1:1 electrolyte corresponding to an ionic strength of 0.15, which served as a canonical baseline condition for modeling the protein–nucleic acid system. All complexes were solvated in an isotropic TIP3P water box^24^, and non-periodic boundary conditions were applied with a 1.0 nm boundary distance. Energy minimization was performed prior to productive simulations, and three independent replicates were run for each complex using the Langevin integrator^23^. A 1.0 nm cutoff was applied to non-bonded interactions for calculation of long-range electrostatic contributions. Constraints were applied to all hydrogen-involving bonds. The system temperature was maintained at 310 K, with temperature fluctuations monitored continuously. The total simulation duration was 100 ns with a 2 fs timestep. System properties including potential energy (kJ/mol), kinetic energy (kJ/mol), and temperature (K) were recorded routinely during sampling. Root-mean-square deviation (RMSD) of non-hydrogen atoms across trajectories was calculated using VMD (v1.9.4a53)^25^, with all structures aligned to the initial simulation frame.

### Validation of the one-pot RPA/CRISPR-Cas12a assays with simulated soil samples

A double-blind method was employed for the evaluation of simulated samples. Concentration gradients of heat-inactivated *B. pseudomallei* (Bp) at 2×10^4^ and 2×10^3^ CFU/mL were established, and heat-inactivated *B. mallei* (Bm, 2×10^6^ CFU/mL), a closely related species, was included as an interference group, while ddH2O was designated as the NTC group. A total of 12 simulated soil samples were prepared. After processing these samples using a thermal lysis-based nucleic acid extraction-free strategy, the RPA/CRISPR one-pot detection method specific for *B. pseudomallei* was applied for evaluation, and a real-time PCR assay for *B. pseudomallei* was concurrently performed for comparative detection (Figure 7B). In addition, control groups consisting of heat-inactivated *B. pseudomallei* diluted in PBS at the same concentrations (2×10^4^ and 2×10^3^ CFU/mL, without soil matrix) were prepared.

The detailed procedures were as follows: The collected soil was resuspended in sterile 1×PBS buffer (approximately 80 mg soil in 500 μL sterile 1×PBS), and heat-inactivated *B. pseudomallei* culture at the corresponding concentration was spiked into the suspension. The mixture was then heated in a metal bath at 99°C for 10 min for thermal lysis. Afterwards, the mixture was allowed to stand to achieve natural sedimentation of soil solids and solid-liquid separation, and the supernatant was aspirated for subsequent detecting experiments.

## Results and Discussion

### Inapplicability of the Established One-pot RPA/CRISPR-Cas12a System to Other Pathogen Targets

Although we referred to the one-pot method study published by Ma et al. ^19^, we still carried out relatively complex parameter optimization—including the concentrations of various components such as Cas12a, crRNA, and ssDNA reporter, as well as the reaction temperature (Figures S1A–S1D), before we could establish the RPA/CRISPR-Cas12a one-pot detection system for *B. anthracis*, achieving a sensitivity of 10 copies/μL (Figures S1E–S1F). However, when this established reaction system was directly applied to *Brucella* and *B. pseudomallei*, no positive signal was observed for either bacterial target (Figure S2). This finding suggests that, in the absence of a generalizable strategy, establishing a one-pot detection method would require individual optimization for each target, resulting in considerable material and time costs as well as an increased risk of failure. Therefore, more broadly applicable strategy is needed to facilitate the development of one-pot RPA/CRISPR-Cas12a detection methods.

### Proposal and Validation of a Universal Base-Mismatch crRNA Strategy

A base-mismatched crRNA strategy is proposed here (Figures 2A and 3A). First, for the crRNAs of *B. pseudomallei* and *Brucella*—namely, *LC1*-crRNA and omp2a-crRNA—single-base mutations were introduced into the seed-region bases to generate mismatches with the target DNA sequences; the corresponding mutation rules are described in the Methods section. Twelve base-mismatched crRNAs for *LC1* and twelve for omp2a were designed separately. The designed base-mismatched crRNAs were then introduced into the established one-pot detection system, target DNA at the same concentration was used, and the effects were evaluated. The results obtained with the unmodified crRNA and with the base-mismatched crRNAs were compared, as shown in Figures 2 and 3.

**Figure 3.**
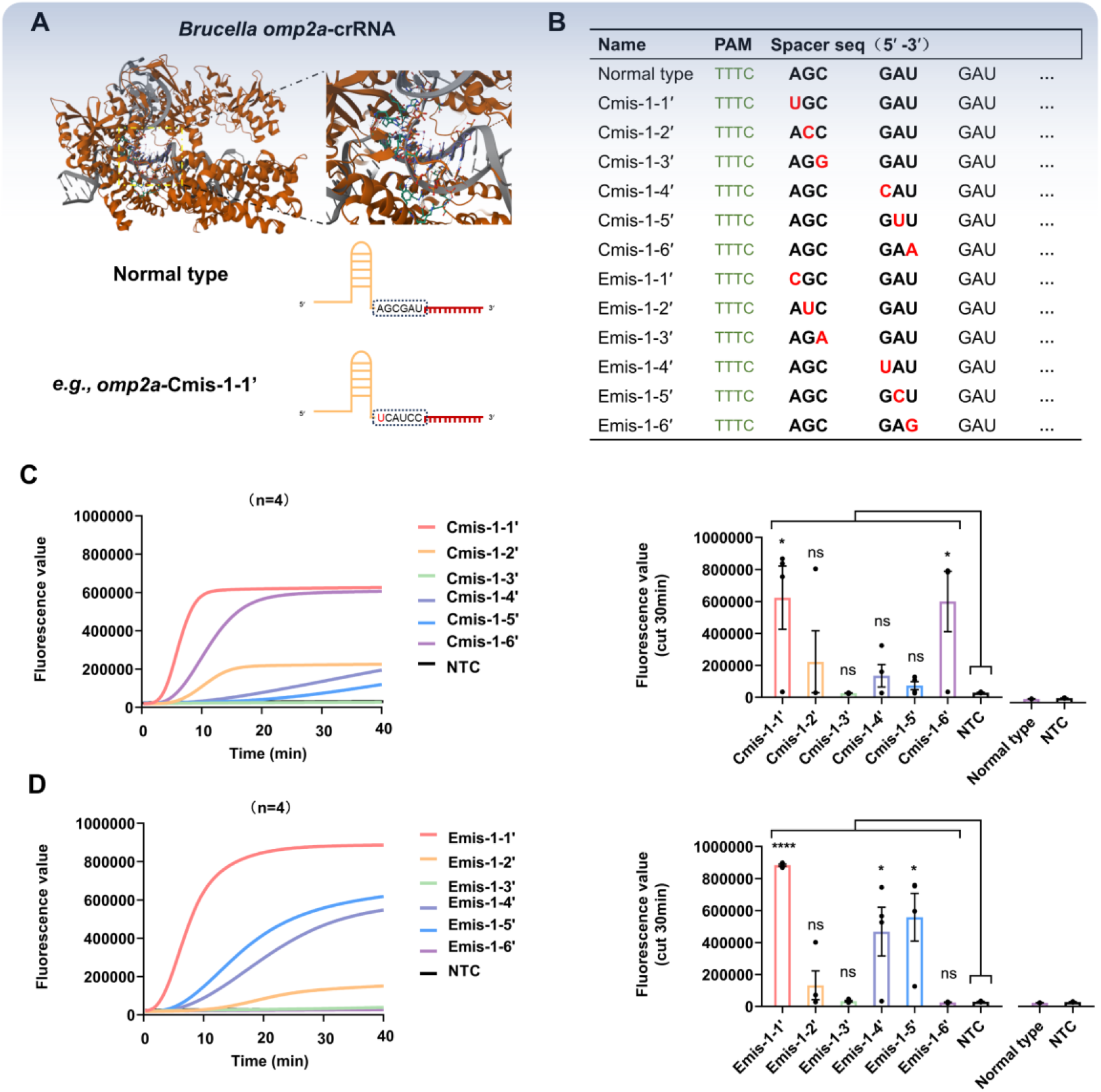
Screening results of base-mismatched crRNAs targeting *Brucella omp2a*. **(A)** Schematic illustration. **(B)** Mutation sites within the seed region of the 12 base-mismatched crRNAs. **(C)** Left panel: fluorescence amplification curves obtained from the Cmis validation group; right panel: bar chart showing the fluorescence signal values measured at the 30-min reaction time point. **(D)** Left panel: fluorescence amplification curves from the Emis validation group; right panel: bar chart of the fluorescence signal values at 30 min. (n=4, mean ±SEM; statistical significance was assessed by t-test, with **** indicating *p* < 0.0001 and ns denoting not significant).

Evaluation results for *B. pseudomallei*-*LC1* are shown in Figure 2. After base mutations were introduced into the seed region, significant positive signals were obtained for *LC1*-Cmis-1-4′ and *LC1*-Emis-1-1′, which were clearly superior to those of the normal-type crRNA. In Figure 3, the evaluation results for *Brucella*-*omp2a* are presented; similarly, following the seed-region base mismatches, *omp2a*-Emis-1-1′ also exhibited marked positive outcomes, and these were notably better than those of the normal-type crRNA. These observations collectively indicate that the designed mismatched crRNAs effectively enhance detection performance for both targets, supporting the applicability of the proposed strategy under the conditions tested.

However, these results were obtained only after crRNA design and wet-lab validation were performed; more intuitive and quantifiable data are therefore needed to further characterize this intriguing phenomenon.

### MD Simulation Analysis of Base-Mismatch crRNA

Based on the wet-lab validation results, several groups of base-mismatched crRNAs with clearly contrasting outcomes were selected from the *B. pseudomallei* and *Brucella* datasets, including unmodified crRNA, *LC1*-Cmis-1-1′, *LC1*-Cmis-1-2′, *LC1*-Cmis-1-4′, *LC1*-Cmis-1-6′, *LC1*-Emis-1-1′, *LC1*-Emis-1-2′, *LC1*-Emis-1-6′, *omp2a*-Cmis-1-1′, *omp2a*-Cmis-1-3′, *omp2a*-Cmis-1-6′, *omp2a*-Emis-1-1′, *omp2a*-Emis-1-3′, and *omp2a*-Emis-1-6′, and these were subjected to MD simulations (Figure 4A). Simulations were performed in triplicate, each for 100 ns.

**Figure 4.**
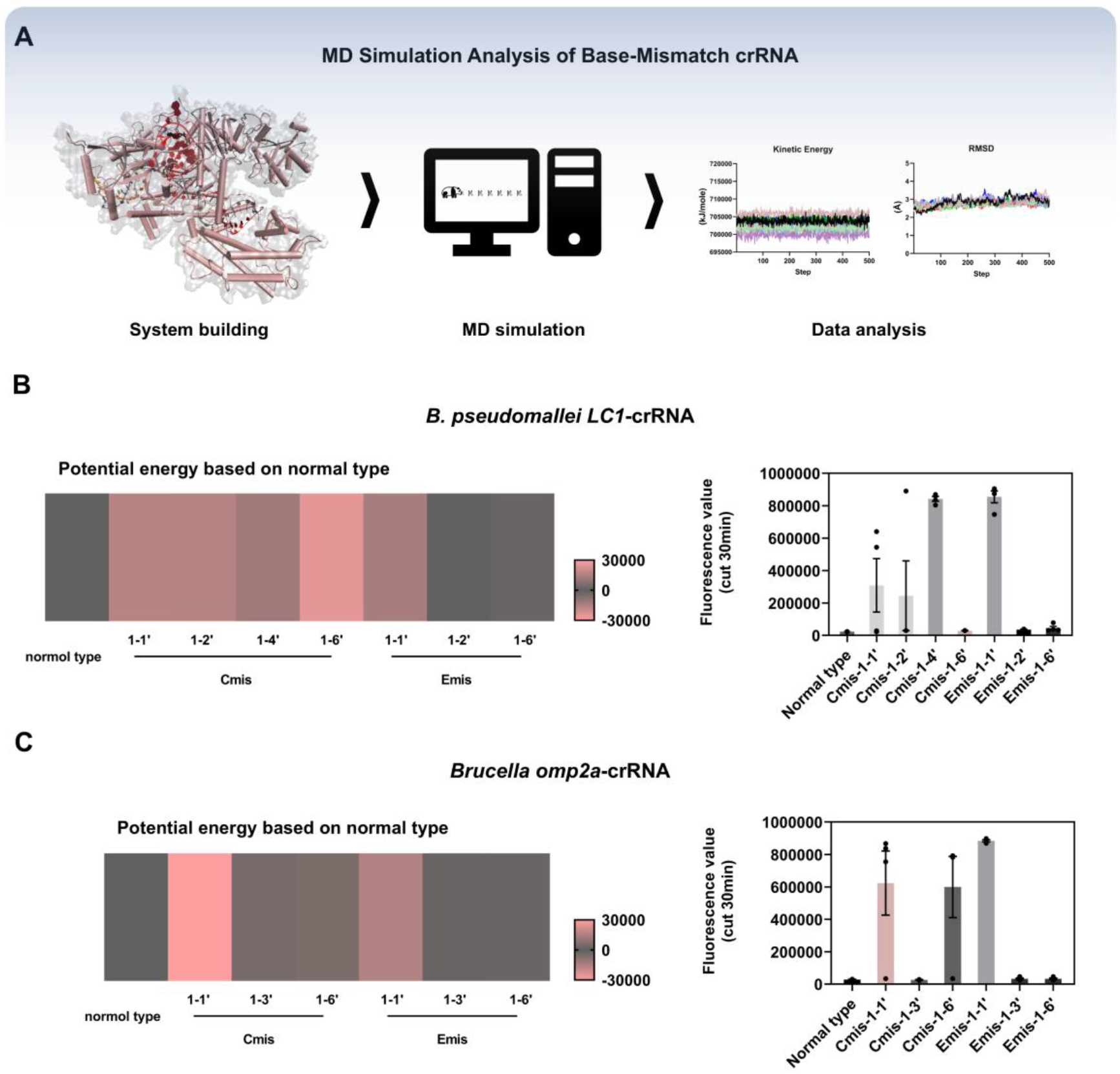
Results of MD simulations. **(A)** Schematic illustration of the simulation approach. **(B)** Simulated PE values obtained for *B. pseudomallei LC1*-crRNA (in kJ/mol). The normal type was assigned a baseline value of 0, and the following mutant values were derived: Cmis1-1’ (–16502.2), Cmis1-2’ (16199.22), Cmis1-4’ (11947.43), Cmis1-6’ (25041.69), Emis1-1’ (13384.41), Emis1-2’ (–160.007), and Emis1-6’ (– 2248.92). **(C)** Simulated PE values for *Brucella omp2a*-crRNA (in kJ/mol), with the normal type set to 0, and mutant values recorded as follows: Cmis1-1’ (–29446.2), Cmis1-3’ (–4710.08), Cmis1-6’ (5226.44), Emis1-1’ (–15517.8), mis1-3’ (–1838.36), and Emis1-6’ (1799.35).

Equilibration of the MD simulations was confirmed by kinetic energy, temperature, and root mean square deviation (RMSD) trajectories, as presented in Figure S3. Following this validation, potential energy (PE) data from each experimental group were compiled and normalized against the unmodified crRNA group, which was defined as the baseline (zero). PE variation for distinct mutation types and sites was then systematically analyzed, and the resulting values were visualized as a color-coded heat map in Figure 4. In this heat map, color intensity was designed to gradually lighten as the magnitude of PE variation increased, ultimately fading to pink at the largest deviations.

Examination of the heat map revealed three distinct patterns of PE variation relative to the unmodified crRNA (Figures 4B and 4C). First, four mutants—*LC1*-Cmis-1-6′, *LC1*-Cmis-1-1′, *LC1*-Cmis-1-2′ and *omp2a*-Cmis-1-1′—exhibited substantially larger PE deviations. These pronounced changes suggested considerable structural or energetic perturbation, which was associated with poor screening performance in subsequent assays. Second, six other mutants—*LC1*-Emis-1-2′, *LC1*-Emis-1-6′, *omp2a*-Cmis-1-3′, *omp2a*-Cmis-1-6′, *omp2a*-Emis-1-3′, and *omp2a*-Emis-1-6′—showed only marginal PE changes, indicating minimal energetic impact; however, this group also yielded unsatisfactory screening results. Notably, a third category, characterized by moderate PE fluctuations, was observed for *LC1*-Cmis-1-4′, *LC1*-Emis-1-1′, and *omp2a*-Emis-1-1′. These intermediate deviations appeared to correspond to the crRNAs that displayed the most pronounced base-mismatch discrimination efficacy in the wet-lab experiments.

Collectively, these findings, together with a mechanistic inference based on the literature, suggest a non-monotonic relationship between PE variation (relative to the normal-type crRNA) and single-tube detection performance. Mechanistically, mutational perturbations of the ternary complex can be rationalized based on established Cas12a principles: crRNA-target hybridization drives conformational checkpoints and RuvC activation, coupling cis target cleavage to trans nonspecific ssDNA cleavage ^26, 27^. Minimal structural perturbation would preserve near-wild-type (normal-type) R-loop propagation and ordered target/non-target cleavage, potentially depleting target DNA before amplification. Severe perturbation, however, impair post-nontarget-strand rearrangements, including PAM-distal unwinding, target-strand delivery to the RuvC site, or R-loop reversibility, lowering both cis and trans cleavage activities^28, 29^. Moderate perturbations that retain seed-to-full R-loop formation and coordinated REC/RuvC docking but limit complete second-strand delivery preserve the trans readout while attenuating cis-cleavage enough to maintain amplifiable target DNA ^30, 31^. PE variation -based screening may therefore help identify crRNA mutants that balancing these two activities in a single tube without wet-lab experiments. Still, the magnitude of variation should be interpreted with caution and confined to a defined range.

### Validation of Sensitivity and Specificity for a One-pot Assay

The selected base-mismatched crRNAs, *LC1*-Cmis-1-4′ and *omp2a*-Emis-1-1′, were used to establish one-pot RPA/CRISPR-Cas12a assays for *B. pseudomallei* and *Brucella*, respectively, and the sensitivity of these assays was subsequently evaluated. Sensitivity evaluation results are presented in Figure 5, both one-pot assays achieved a detection limit of 10 copies/μL, consistent with that obtained for the *B. anthracis* assay.

**Figure 5.**
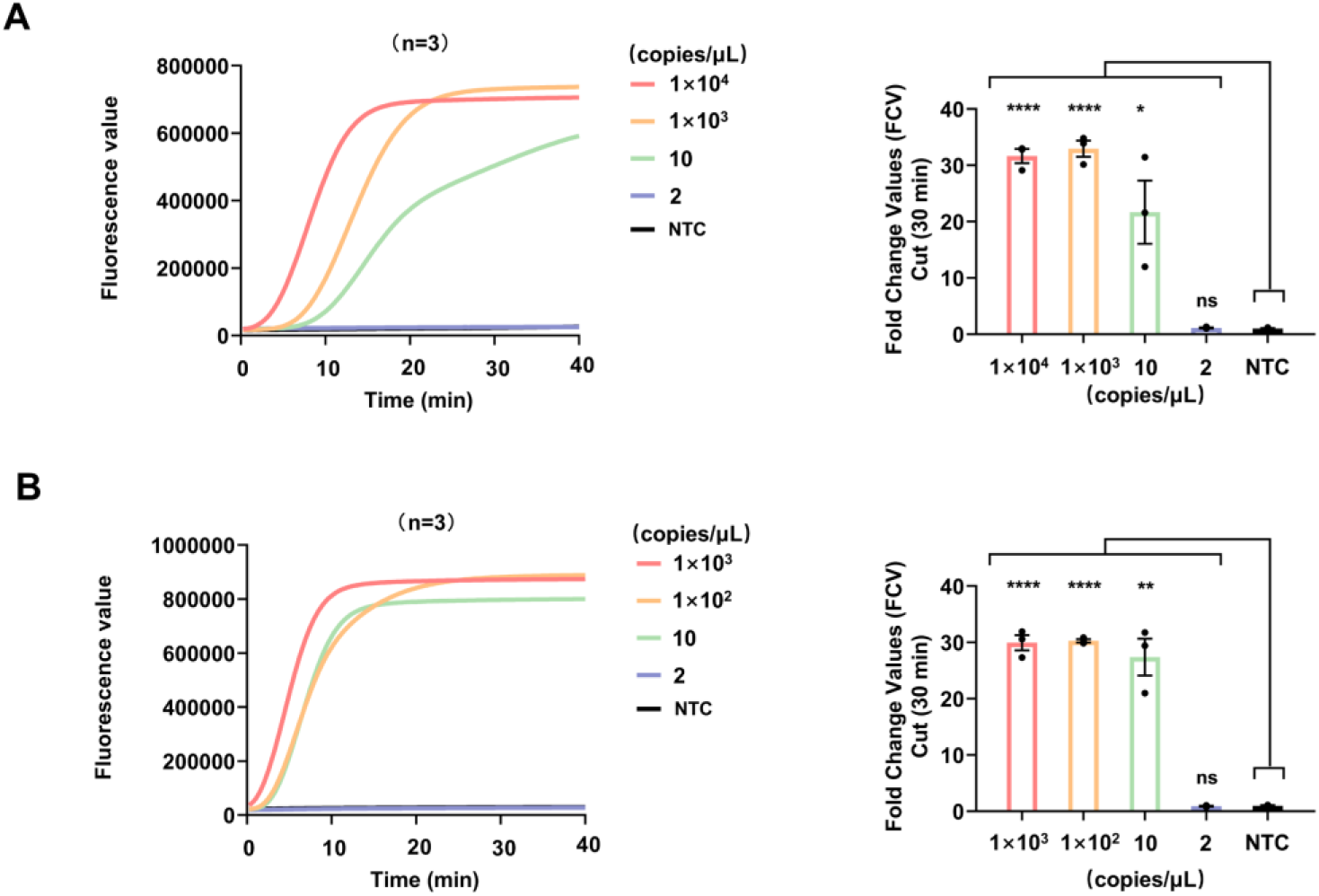
Evaluation of the sensitivity of the one-pot RPA/CRISPR assay. **(A)** *B. pseudomallei*: left panel, fluorescence amplification curves; right panel, bar chart of fluorescence signal values measured at the 30 min reaction time point (n=3, mean±SEM). Statistical significance was assessed by t-test, with the following results for each concentration group: 1×10^4^ copies/μL (****, *p*<0.0001), 1×10^3^ copies/μL (****, *p*<0.0001), 10 copies/μL (*, *p*=0.0212), and 2 copies/μL (ns, not significant). **(B)** *Brucella:* left panel, fluorescence amplification curves; right panel, bar chart of signal values at 30 min (n = 3, mean ± SEM). t-test was performed, yielding: 1×10^3^ copies/μL (****, *p*<0.0001), 1×10^2^ copies/μL (****, *p*<0.0001), 10 copies/μL (**, *p*=0.0013), and 2 copies/μL (ns, not significant).

To evaluate the specificity of the one-pot assays for the three bacterial species, non-target bacterial mixed genomic DNA (100 pg/μL) and mixed genomic DNA derived from human whole blood and soil samples (1 ng/μL) were used as interference groups. In parallel, positive control (PC; target bacterial genomic DNA, 100 fg/μL) and no-template control (NTC) groups were also established. Evaluation results are presented in Figure 6. For each bacterial one-pot assay, positive signals were detected exclusively in the PC group, whereas no nonspecific amplification was observed in the interference groups or the NTC group, indicating satisfactory specificity for all assays. Moreover, these results suggest that the single-base mismatch strategy of crRNA in the seed region does not trigger Cas12a off-target effects within the RPA-CRISPR detection system, which would produce false-positive signals in CRISPR-based in vitro diagnostics.

**Figure 6.**
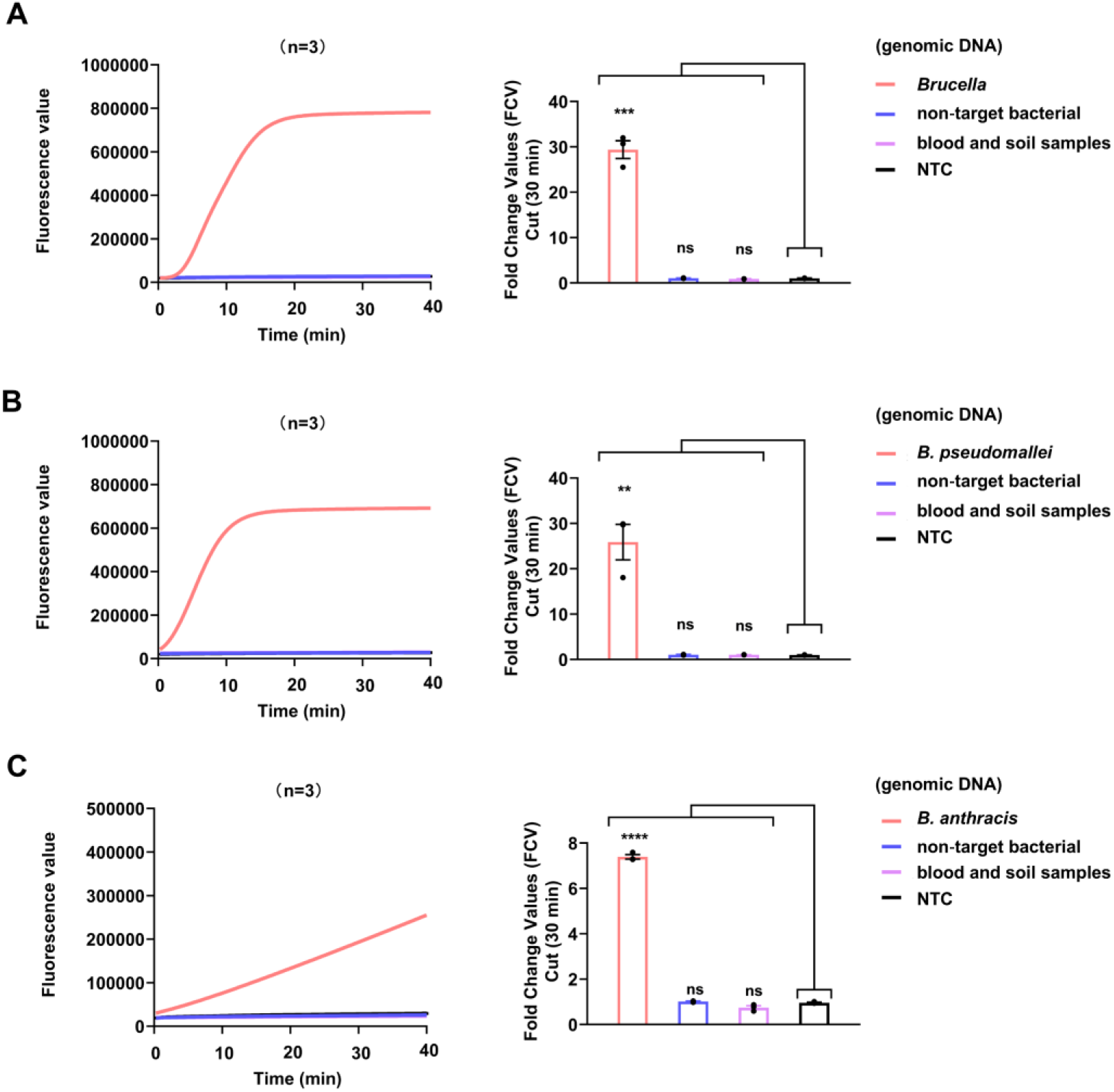
Specificity evaluation results. Fluorescence signal values measured at the 30 min reaction time point are presented as bar charts (n=3, mean±SEM). Statistical comparisons were performed using the t-test. **(A)** *Brucella*, ***, *p*=0.0001. **(B)** *B. pseudomallei*, **, *p*=0.0031. **(C)** *B. anthracis*, ****, *p*<0.0001.

### On-Site Independent Work Unit Setup and Evaluation

The one-pot RPA/CRISPR-Cas12a detection systems for the three bacterial targets were prepared into lyophilized pellets via a freeze-drying method by Tolo Biotech Co.,Ltd (Wuxi, Jiangsu, China), and these pellets were subsequently loaded into 200 μL transparent PCR centrifuge tubes. Fluorescence signal collection during the detection process was performed using a battery-powered portable nucleic acid isothermal amplification instrument (Genie II), enabling real-time monitoring of signals on the display screen. Together with other necessary consumables, this configuration constituted an independent work unit (Figure 7A) suitable for potential field deployment.

**Figure 7.**
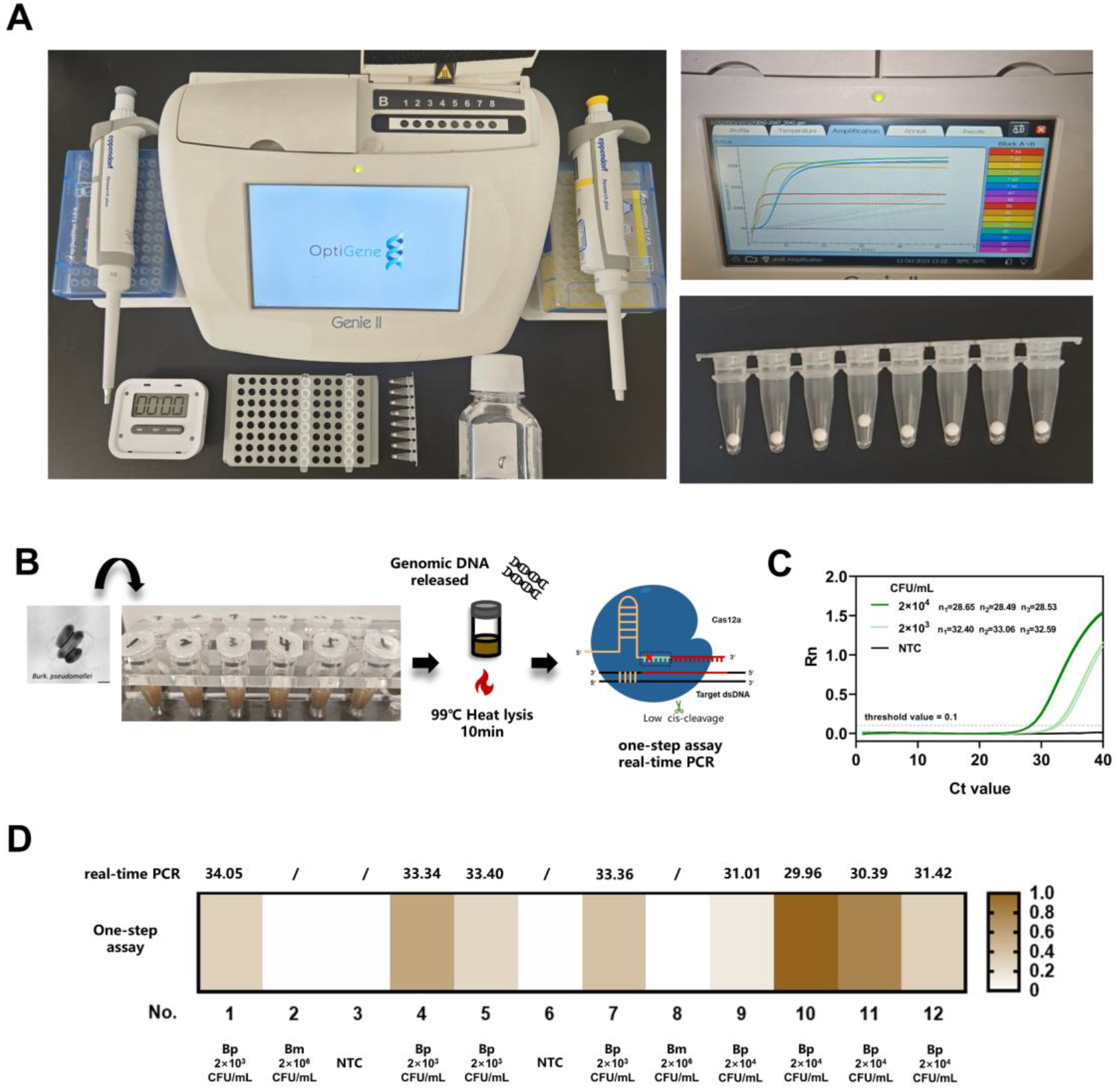
(A) An independent work unit suitable for on-site operations was established, and a photograph of the one-pot RPA/CRISPR lyophilized microspheres is presented. A flowchart of the extraction-free nucleic acid detection procedure for *B. pseudomallei* in soil-simulated samples is shown. (C) Real-time PCR results obtained from the PBS dilution control group are displayed. (D) A heat map of the extraction-free nucleic acid detection results of *B. pseudomallei* from the soil-simulated samples is provided.

In addition, the prepared lyophilized pellets for one-pot detection of the three targets were stored at room temperature for 10 and 30 days. Recombinant positive plasmids of *B. anthracis, B. pseudomallei*, and *Brucella* at concentrations of 100 copies/μL and 10 copies/μL were then used as template DNA, and ddH2O served as the NTC group, with two replicates per group. Subsequently, 10 μL of resuspension buffer (9 μL ddH2O + 1 μL MgOAc) and 10 μL of template DNA were added to reconstitute the lyophilized microspheres, and the reaction was performed at 39 °C in the Genie ? instrument. As shown in Figure S4, after storage at room temperature for 10 and 30 days, recombinant positive plasmids at 100 copiesrμL were consistently detected, suggesting that the system was largely capable of meeting the demands of rapid on-site screening. Nevertheless, further optimization of the lyophilization process and formulation remains necessary to more fully approximate the sensitivity and efficiency attainable with the liquid-phase assay.

This prepared lyophilized pellets for one-pot detection was validated with simulated soil samples. Following identical extraction-free processing (99Ċ, 10 min), these PBS controls were also detected by the real-time PCR method. The results obtained from three replicates are shown in Figure 7C, with Ct values of approximately 28.56 (2×10^4^ CFU/mL) and 32.68 (2×10^3^ CFU/mL). However, simulated soil samples with an average Ct value of approximately 30.70 (2×10^4^ CFU/mL) and 33.54 (2×10^3^ CFU/mL) in Figure 7D, suggesting that the soil matrix may have interfered with both the lysis efficiency of the extraction-free strategy and the detection performance of the real-time PCR assay. This observation underscores the importance of cautious interpretation of diagnostic results derived from complex environmental matrices and highlights potential challenges associated with applying simplified nucleic acid preparation protocols directly to field samples without prior purification.

Both the RPA/CRISPR-Cas12a one-pot detection method and the real-time PCR method were able to detect *B. pseudomallei* in the simulated soil samples, with no apparent interference observed from *B. mallei* or from solid inorganic and organic substances inherently present in the soil (Figure 7D). These results indicate that the developed approach possesses adequate specificity and robustness under the conditions tested, supporting its potential suitability for diagnostic applications involving complex sample matrices.

## Conclusion

This study establishes a one-pot RPA/CRISPR-Cas12a diagnostic platform by introducing base mismatches into the seed region (positions 1–6) of crRNA. This strategy effectively suppresses the cis-cleavage activity of Cas12a, thereby allowing nucleic acid amplification and CRISPR detection to proceed within a single tube without the need for costly chemically modified crRNAs or laborious target-specific PAM rescreening. Integrating MD simulations with wet-lab validation, we found a non-monotonic relationship between PE variation (relative to normal-type crRNA) and single-tube Cas12a detection: both too small and too large PE deviations failed, whereas moderate perturbations performed best. This observation offers a potentially useful framework to guide future crRNA design and may help reduce experimental trial-and-error, pending further validation across additional targets. Furthermore, successful lyophilization of the components of this one-pot reaction system into portable, battery-operated work units, together with robust performance in simulated soil samples, suggests the method’s potential practical value for field deployment in resource-limited settings.

Nevertheless, certain challenges remain. Future work will focus on improving lyophilization processes for higher sensitivity and longer RT storage time; incorporating more efficient nucleic acid release strategies, and expanding the high-risk pathogens panel to further evaluate the field-deployment potential of this diagnostic platform for biothreats or infectious disease surveillance.

## Authors contributions

Conceptualization: Y.Y.; Data curation: Y.Y., X.J.H.; Methodology: YY, X.J.H., W.X., H.M.D; Investigation: X.J.H., Y.B.; Visualization: X.J.H, W.X.; Funding acquisition: Y.Y., X.W.W.; Project administration: Y.Y., R.H.G., W.J.L.; Supervision: W.J.L., R.H.G.; Writing – original draft: X.J.H.; Writing – review & editing: Y.Y., R.H.G.

## Data availability statement

The data that support the findings of this study are available from the corresponding author upon reasonable request.

## Acknowledgments

No funding was received for this research.

## Reference

1. L. Chen, M.L. Hu, X.M. Zhou, Trends in developing one-pot CRISPR diagnostics strategies, Trends Biotechnol. 2025, 43, 98–110.

2. Z.J. Tu, Y.X. Wang, X. Qian, J.L. Chen, L.Y. Li, T. Wang, H. Chen, H. Wei, P. Yang, J. Pan, S. Wang, Z. Rong, Plasmonic magnetic nanoparticles-enabled universal enrichment, photothermal lysis, and duplex CRISPR detection of bacteria in urine samples, Biosens. Bioelectron. 2026, 302, 118518.

3. Y.Y. Yu, M.R. Jin, W.G. Yuan, Y.J. Gong, S.W. Li, X. Qin, J. Hou, J. Liu, S. Liu, H. Li, Y. Chu, Y. Wang, Y. Zhang, F. Fang, W. Hao, Y. Gu, Q. Fan, J. Lin, D. Pang, X. Zhang, Engineered crRNA Drives RPA-T7-CRISPR/Cas14a Cascade for Ultrasensitive Detection of ctDNA PIK3CA H1047R, Adv. Sci. 2025, 12, e07126.

4. N. Kim, D.S. Collins, N.M. Donghia, B.S. Miller, H.M. Sallum, S.R. Lybbert, E. Perini, J.B. Niemi, J.J. Collins, P.Q. Nguyen, A field-deployable CRISPR-based biosensing platform for monitoring marine ecosystems, Nat. Sustain. 2026, 9, 51–64.

5. J.C. Rolando, A. Thieme, N.E. Weckman, N. Kim, H. de Puig, X. Tan, E. Cotnoir, V. Chaturvedi, J.J. Collins, D.R. Walt, Digital CRISPR-based diagnostics for quantification of Candida auris and resistance mutations, Nat. Biomed. Eng. 2026, 9, 51–64.

6. H.Y. Liu, H. Yin, L.S. Xiu, W.Y. Wu, Q.Q. Hu, Y.C. Xia, B. Garcia, S. Shifa, H. Chen, M. Li, K. Yin, One-Pot Isothermal Nucleic Acid Amplification Assisted CRISPR/Cas Detection Technology: Challenges, Strategies, and Perspectives, Adv. Sci. 2025, 12, e06716.

7. X.H. Tong, K. Zhang, Y. Han, T.L. Li, M. Duan, R.J. Ji, X.G. Wang, X. Zhou, Y. Zhang, H. Yin, Fast and sensitive CRISPR detection by minimized interference of target amplification, Nat. Chem. Biol. 2024, 20, 885–893.

8. S.H. Lu, X.H. Tong, Y. Han, K. Zhang, Y.Z. Zhang, Q.B Chen, J.Y. Duan X.L. Lei, M.H. Huang, Y. Qiu, D.Y. Zhang, X. Zhou, Y. Zhang, H. Yin, Fast and sensitive detection of SARS-CoV-2 RNA using suboptimal protospacer adjacent motifs for Cas12a, Nat. Biomed. Eng. 2022, 6, 286–297.

9. L. Yang, G.W. Chen, J. Wu, W. Wei, C. Peng, L. Ding, X.Y. Chen, X. Xu, X. Wang, J. Xu, A PAM-Free One-pot Asymmetric RPA and CRISPR/Cas12b Combined Assay (OAR-CRISPR) for Rapid and Ultrasensitive DNA Detection, Anal. Chem. 2024, 96, 5471–5477.

10. Z.H. Cheng, X.Y. Luo, S.S. Yu, D. Min, S.X. Zhang, X.F. Li, J.J. Chen, D.F. Liu, H.Q. Yu, Tunable control of Cas12 activity promotes universal and fast one-pot nucleic acid detection, Nat. Commun. 2025, 16, 1166.

11. T. Tian, H.R Xiao, X.Y. Guo, Y.X. Chen, Z.Q. Qiu, T. Zhang, M.Y. Chen, W.W. Qi, P.G. Cai, M. Cheng, X.M. Zhou, Identification of a key nucleotide influencing Cas12a crRNA activity for universal photo-controlled CRISPR diagnostics, Nat. Commun. 2025, 16, 6694.

12. M.L. Hu, Z.Q. Qiu, Z.R. Bi, T. Tian, Y.Z. Jiang, X.M. Zhou, Photocontrolled crRNA activation enables robust CRISPR-Cas12a diagnostics, PNAS. 2022, 119, e2202034119.

13. M. Cheng, Y. Wang, W. Lin, J. Ye, M. Wu, B. Xiang, L. Liu, B. Sun, A Universal Light-Activated CRISPR-RNA Based on Split Direct Repeat for One-Pot Cas12a Nucleic Acid Detection, Anal. Chem. 2026, 98, 706–716.

14. L. Villiger, J. Joung, L. Koblan, J. Weissman, O.O. Abudayyeh, J.S. Gootenberg, CRISPR technologies for genome, epigenome and transcriptome editing, Nat. Rev. Mol. Cell Biol. 2024, 25, 464–487.

15. C. Guo, S. Zhang, R. Yerramsetti, J.Y. Zhang, X. Guan, R. Yang, C.Y. Hou, M. Pei, K.T. Schalper, X. Liu, Z. Li, L. Perpetua, W.Q. Gan, O. Ibrahim, R.A. Clark, C.C. Liu, Single-nucleotide variant profiling in liquid biopsy with RECO-Cas, Sci. Adv. 2026, 12, eaed1757.

16. H.K. Kim, M. Song, J. Lee, M. A Vipin, J. Soobin, K. Young-Mook, C. Jae W W. Euijeon, H.C Koh, J.W. Nam, H. Kim, In vivo high-throughput profiling of CRISPR-Cpf1 activity, Nat. Methods. 2016, 14, 153–159.

17. Z.H. Yao, W.L. Li, K.Y. He, H.M. Wang, Y. Xu, Q. Wu, L. Wang, Y. Nie, Facilitating crRNA Design by Integrating DNA Interaction Features of CRISPR-Cas12a System, Adv. Sci. 2025, 12, e2501269.

18. J.H. Xu, X.R. Bai, X.L.L. Zhang, B. Yuan, L. Lin, Y. Guo, Y.J. Cui, J. Liu, H. Cui, X. Ren, J.L. Wang, Y. Yuan, Development and application of DETECTR-based rapid detection for pathogenic Bacillus anthracis, Anal. Chim. Acta. 2023, 1247, 340891.

19. Y.J. Ma, H.J. Wei, Y.X. Wang, X.D. Cheng, H. Chen, X.S. Yang, H.S. Zhang, Z. Rong, S.Q. Wang, Efficient magnetic enrichment cascade single-step RPA-CRISPR/Cas12a assay for rapid and ultrasensitive detection of Staphylococcus aureus in food samples, J. Hazard. Mater. 2024, 465, 133494.

20. J.X. Zhang, J.H. Xu, B. Yuan, X.D. Wang, X.H. Mao, J.L. Wang, X.L.L. Zhang, Y. Yuan, Detection of Burkholderia pseudomallei with CRISPR-Cas12a based on specific sequence tags, Front. Public. Health. 2023, 11, 1153352.

21. J.H. Xu, J.F. Ma, Y.W. Li, L. Kang, B. Yuan, S.Q. Li, J. Chao, L.H. Wang, J.L. Wang, S. Su, Y. Yuan, A general RPA-CRISPR/Cas12a sensing platform for Brucella spp. detection in blood and milk samples, Sensors and Actuators: B. Chemical 2022, 364, 131864.

22. P. Eastman, R. Galvelis, R.P. Peláez, C.R.A. Abreu, S.E. Farr, E. Gallicchio, A. Gorenko, M.M. Henry, F. Hu, J. Huang, A. Krämer, J. Michel, J.A. Mitchell, V.S. Pande, J.P. Rodrigues, J. Rodriguez-Guerra, A.C. Simmonett, S. Singh, J. Swails, P. Turner, Y. Wang, I. Zhang, J.D. Chodera, G. Fabritiis, T.E. Markland, OpenMM 8: Molecular Dynamics Simulation with Machine Learning Potentials, J Phys Chem B. 2024, 128, 109–116.

23. R. Salomon-Ferrer, A.W. Götz, D. Poole, S. Grand, R.C. Walker, Routine Microsecond Molecular Dynamics Simulations with AMBER on GPUs. 2. Explicit Solvent Particle Mesh Ewald, J Chem Theory Comput. 2013, 9, 3878–88.

24. D. Nayar, M. Agarwal, C. Chakravarty, Comparison of Tetrahedral Order, Liquid State Anomalies, and Hydration Behavior of mTIP3P and TIP4P Water Models, J Chem Theory Compu. 2011, 7, 3354–67.

25. Humphrey W, Dalke A, S. K, VMD: visual molecular dynamics, J Mol Graph. 1996, 14, 33-8, 27–8.

26. S. Stella, P. Mesa, J. Thomsen, B. Paul, P. Alcón, S. Jensen, B. Saligram, M. Moses, N. Hatzakis, G. Montoya, Conformational Activation Promotes CRISPR-Cas12a Catalysis and Resetting of the Endonuclease Activity, Cell. 2018, 175, 1856–1871.e21.

27. D.C. Swarts, M. Jinek, Mechanistic Insights into the cis- and trans-Acting DNase Activities of Cas12a, Mol. Cell. 2019, 73, 589–600.e4.

28. H. Son, J. Park, I. Hwang, Y. Jung, S. Bae, S. Lee, Mg2+-dependent conformational rearrangements of CRISPR-Cas12a R-loop complex are mandatory for complete double-stranded DNA cleavage, PNAS. 2021, 118, e2113747118.

29. I. Strohkendl, F.A. Saifuddin, J.R. Rybarski, I.J. Finkelstein, R. Russell, Kinetic Basis for DNA Target Specificity of CRISPR-Cas12a, Mol. Cell. 2018, 71, 816–824.e3.

30. I. Strohkendl, A. Saha, C. Moy, A.-H. Nguyen, M. Ahsan, R. Russell, G. Palermo, D.W. Taylor, Cas12a domain flexibility guides R-loop formation and forces RuvC resetting, Mol. Cell. 2024, 84, 2717–2731.e6.

31. J.C. Cofsky, D. Karandur, C.J. Huang, I.P. Witte, J. Kuriyan, J.A. Doudna, CRISPR-Cas12a exploits R-loop asymmetry to form double-strand breaks, Elife. 2020, 9, e55143

