## Supplementary Material for "Base-mismatched crRNA in CRISPR Enabled Universal One-pot Rapid Nucleic Acid Detection"

**Table S1** The nucleic acid sequences involved in this study

| **Name** | **Sequence（5**′ **-3**′**）** |
| --- | --- |
| *5345*-F^[1]^ | TTGGTAGATCAGTTGGTGGGATAAC |
| *5345*-R | TGAGTTTTACCTTCTGACAACAATATTTC |
| *5345*-crRNA | UAAUUUCUACUAAGUGUAGAUAAUUGUGAAUUUGUCUGGCACAUGG |
| *LC1*-F^[2]^ | CGGGTTTCCAGAAGGGCTGCAAGCACCAAAT |
| *LC1*-R | TGCAATGTCTTACAACAAGCCATGCCGTCATCTTC |
| *LC1*-crRNA | UAAUUUCUACUAAGUGUAGAUGCAUCCGCCGACCGAUUUGAUGUU |
| omp2a-F^[3]^ | AACTCGGCACACTCGGCACCTACACTCAGC |
| omp2a-R | GGATTCGTCGATACCAACCTTGAAACCACCAAGCG |
| omp2a-crRNA | UAAUUUCUACUAAGUGUAGAUAGCGAUGAUCGUGAUGUCGCUGAU |
| crRNA-*LC1*-Cmis(1-1′) | UAAUUUCUACUAAGUGUAGAUCCAUCCGCCGACCGAUUUGAUGUU |
| crRNA-*LC1*-Cmis(1-2′) | UAAUUUCUACUAAGUGUAGAUGGAUCCGCCGACCGAUUUGAUGUU |
| crRNA-*LC1*-Cmis(1-3′) | UAAUUUCUACUAAGUGUAGAUGCUUCCGCCGACCGAUUUGAUGUU |
| crRNA-*LC1*-Cmis(1-4′) | UAAUUUCUACUAAGUGUAGAUGCAACCGCCGACCGAUUUGAUGUU |
| crRNA-*LC1*-Cmis(1-5′) | UAAUUUCUACUAAGUGUAGAUGCAUGCGCCGACCGAUUUGAUGUU |
| crRNA-*LC1*-Cmis(1-6′) | UAAUUUCUACUAAGUGUAGAUGCAUCGGCCGACCGAUUUGAUGUU |
| crRNA-*LC1*-Emis(1-1′) | UAAUUUCUACUAAGUGUAGAUUCAUCCGCCGACCGAUUUGAUGUU |
| crRNA-*LC1*-Emis(1-2′) | UAAUUUCUACUAAGUGUAGAUGAAUCCCCGACCGAUUUGAUGUU |
| crRNA-*LC1*-Emis(1-3′) | UAAUUUCUACUAAGUGUAGAUGCCUCCGCCGACCGAUUUGAUGUU |
| crRNA-*LC1*-Emis(1-4′) | UAAUUUCUACUAAGUGUAGAUGCAGCCGCCGACCGAUUUGAUGUU |
| crRNA-*LC1*-Emis(1-5′) | UAAUUUCUACUAAGUGUAGAUGCAUACGCCGACCGAUUUGAUGUU |
| crRNA-*LC1*-Emis(1-6′) | UAAUUUCUACUAAGUGUAGAUGCAUCAGCCGACCGAUUUGAUGUU |
| crRNA-*omp2a*-Cmis(1-1′) | UAAUUUCUACUAAGUGUAGAUUGCGAUGAUCGUGAUGUCGCUGAU |
| crRNA-*omp2a*-Cmis(1-2′) | UAAUUUCUACUAAGUGUAGAUACCGAUGAUCGUGAUGUCGCUGAU |
| crRNA-*omp2a*-Cmis(1-3′) | UAAUUUCUACUAAGUGUAGAUAGGGAUGAUCGUGAUGUCGCUGAU |
| crRNA-*omp2a*-Cmis(1-4′) | UAAUUUCUACUAAGUGUAGAUAGCCAUGAUCGUGAUGUCGCUGAU |
| crRNA-*omp2a*-Cmis(1-5′) | UAAUUUCUACUAAGUGUAGAUAGCGUUGAUCGUGAUGUCGCUGAU |
| crRNA-*omp2a*-Cmis(1-6′) | UAAUUUCUACUAAGUGUAGAUAGCGAAGAUCGUGAUGUCGCUGAU |
| crRNA-*omp2a*-Emis(1-1′) | UAAUUUCUACUAAGUGUAGAUCGCGAUGAUCGUGAUGUCGCUGAU |
| crRNA-*omp2a*-Emis(1-2′) | UAAUUUCUACUAAGUGUAGAUAUCGAUGAUCGUGAUGUCGCUGAU |
| crRNA-*omp2a*-Emis(1-3′) | UAAUUUCUACUAAGUGUAGAUAGAGAUGAUCGUGAUGUCGCUGAU |
| crRNA-*omp2a*-Emis(1-4′) | UAAUUUCUACUAAGUGUAGAUAGCUAUGAUCGUGAUGUCGCUGAU |
| crRNA-*omp2a*-Emis(1-5′) | UAAUUUCUACUAAGUGUAGAUAGCGCUGAUCGUGAUGUCGCUGAU |
| crRNA-*omp2a*-Emis(1-6′) | UAAUUUCUACUAAGUGUAGAUAGCGAGGAUCGUGAUGUCGCUGAU |
| ssDNA-reporter | FAM-CCCCCC-BHQ1 |

**Table S2** The composition of RPA/CRISPR mixture

| **Name** | **Concentration** | **Volume** |
| --- | --- | --- |
| RPA Forward Primer | 10 μM | 2.5 μL |
| RPA Reverse Primer | 10 μM | 2.5 μL |
| RPA Buffer |  | 46 μL |
| Cas12a | 10 μM | 0.5 μL |
| crRNA | 2 μM | 1.25 μL |
| ssDNA probe | 10 μM | 10 μL |
| RNase inhibitor | 40000 U/mL | 2.5 μL |
| DDH_2_O |  | 4.75 μL |
| Total |  | 70 μL |

The prepared RPA/CRISPR mixture was added to a single-reaction RPA lyophilized pellet, and the mixture was vortexed to ensure thorough mixing. The mixed solution was then aliquoted into five tubes, with 14 μL added to each tube. Subsequently, 5 μL of template DNA and 1 μL of MgOAc were added to each tube, bringing the final reaction volume to 20 μL. After vortexing, the tubes were placed into a Genie-II nucleic acid isothermal amplification system, pre-incubated at 39°C for 10 min, and then incubated at 39°C for 1 h. The fluorescence curves were monitored throughout the reaction.

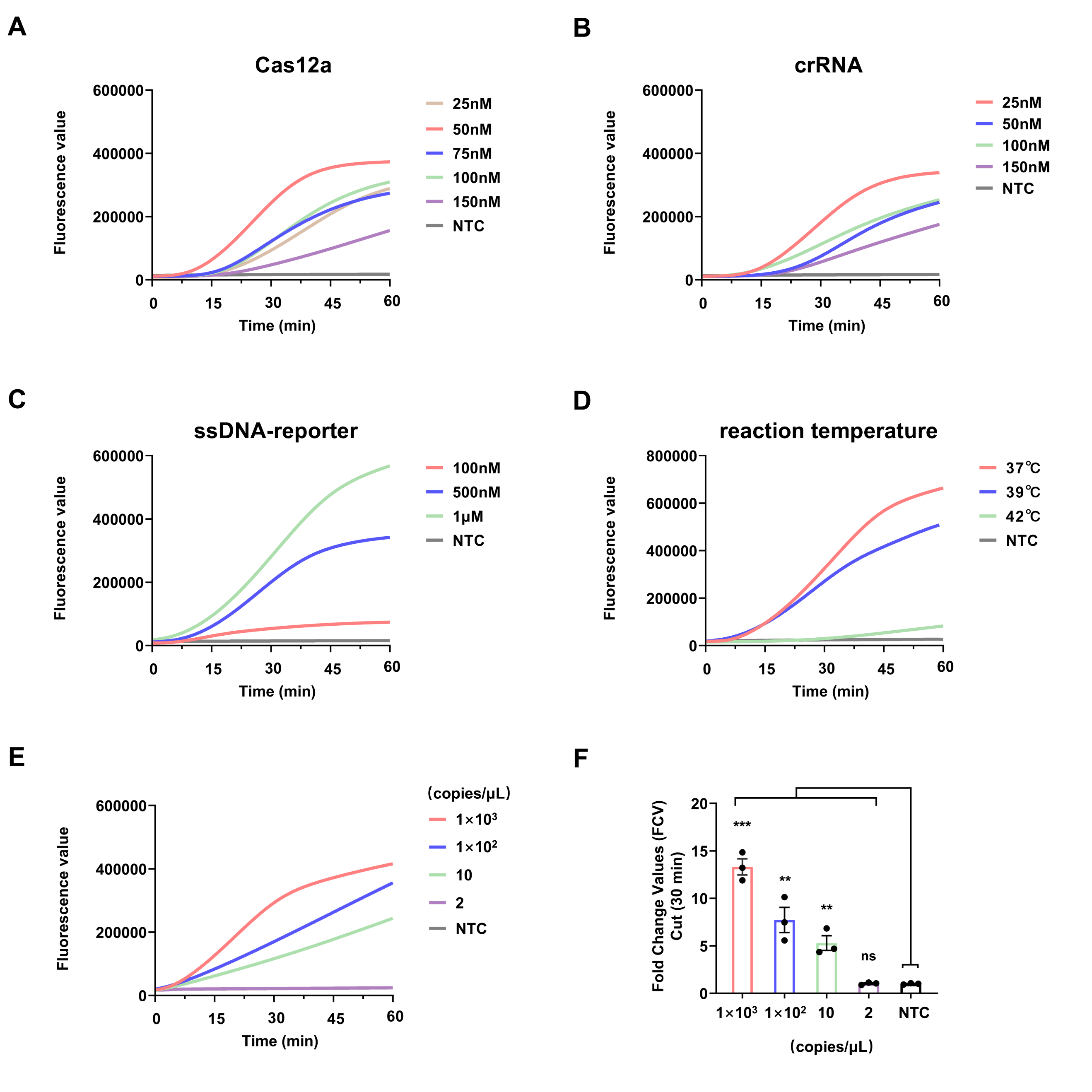

**Figure S1** (A–D) Optimization results for the reaction conditions of the one‑step RPA/CRISPR assay targeting *B. anthracis*. (E, F) Sensitivity evaluation results of the one‑step RPA/CRISPR assay for *B. anthracis*. Fluorescence signal values measured at the 30‑min reaction time point are presented as bar charts (n=3, mean ±SEM). Statistical comparisons were performed using the t‑test, yielding the following significance levels for each concentration group: 1×10³ copies/μL (***, p=0.0001), 1×10² copies/μL (**, p=0.0071), 10 copies/μL (**, p=0.0053), and 2 copies/μL (ns, not significant).

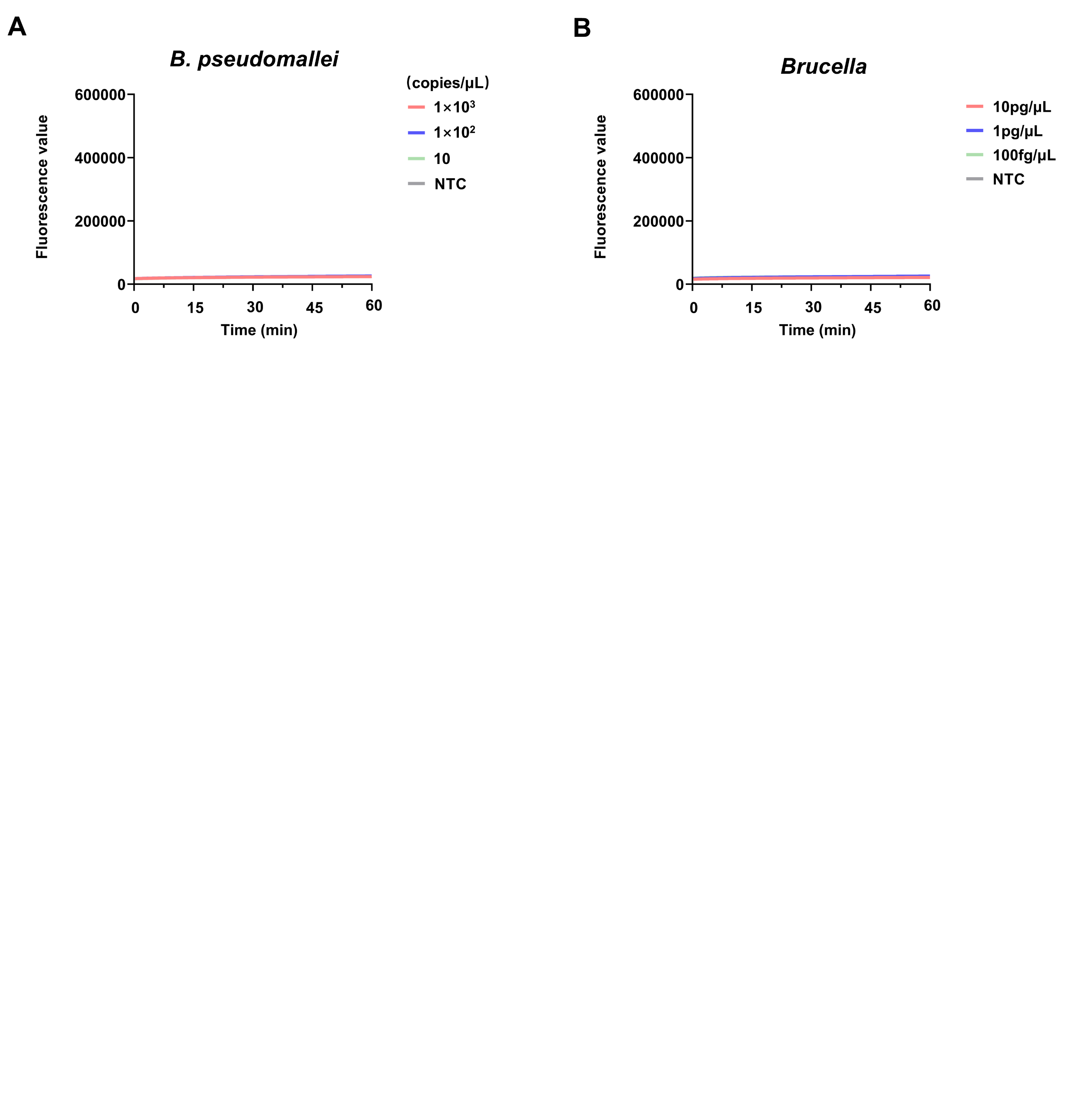

**Figure S2** (A) *B. pseudomallei* and (B) *Brucella* were directly tested using the one‑step RPA/CRISPR detection system originally established for *B. anthracis*.

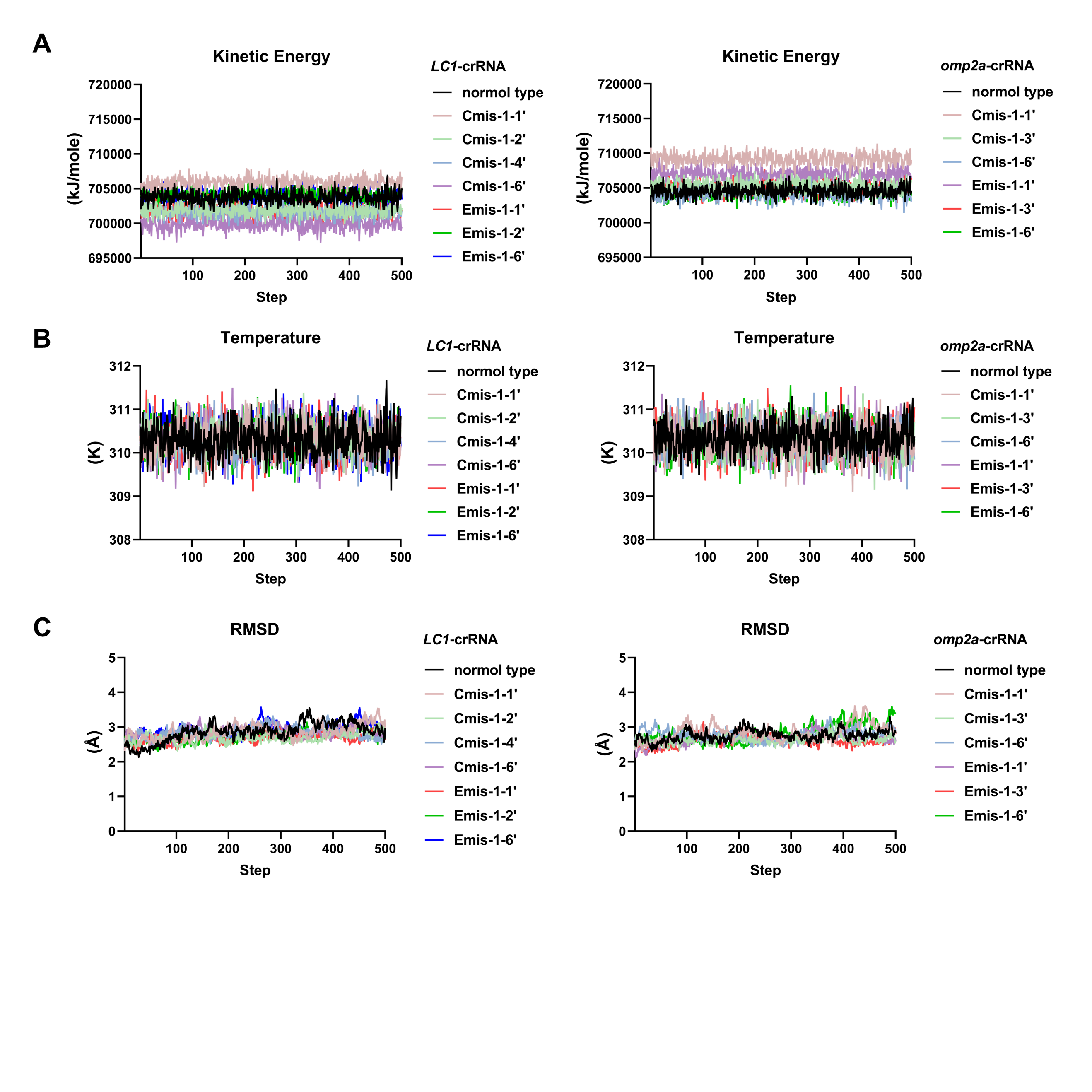

**Figure S3** Molecular dynamics (MD) simulation results: (A) Kinetic Energy, (B) Temperature, (C) RMSD.

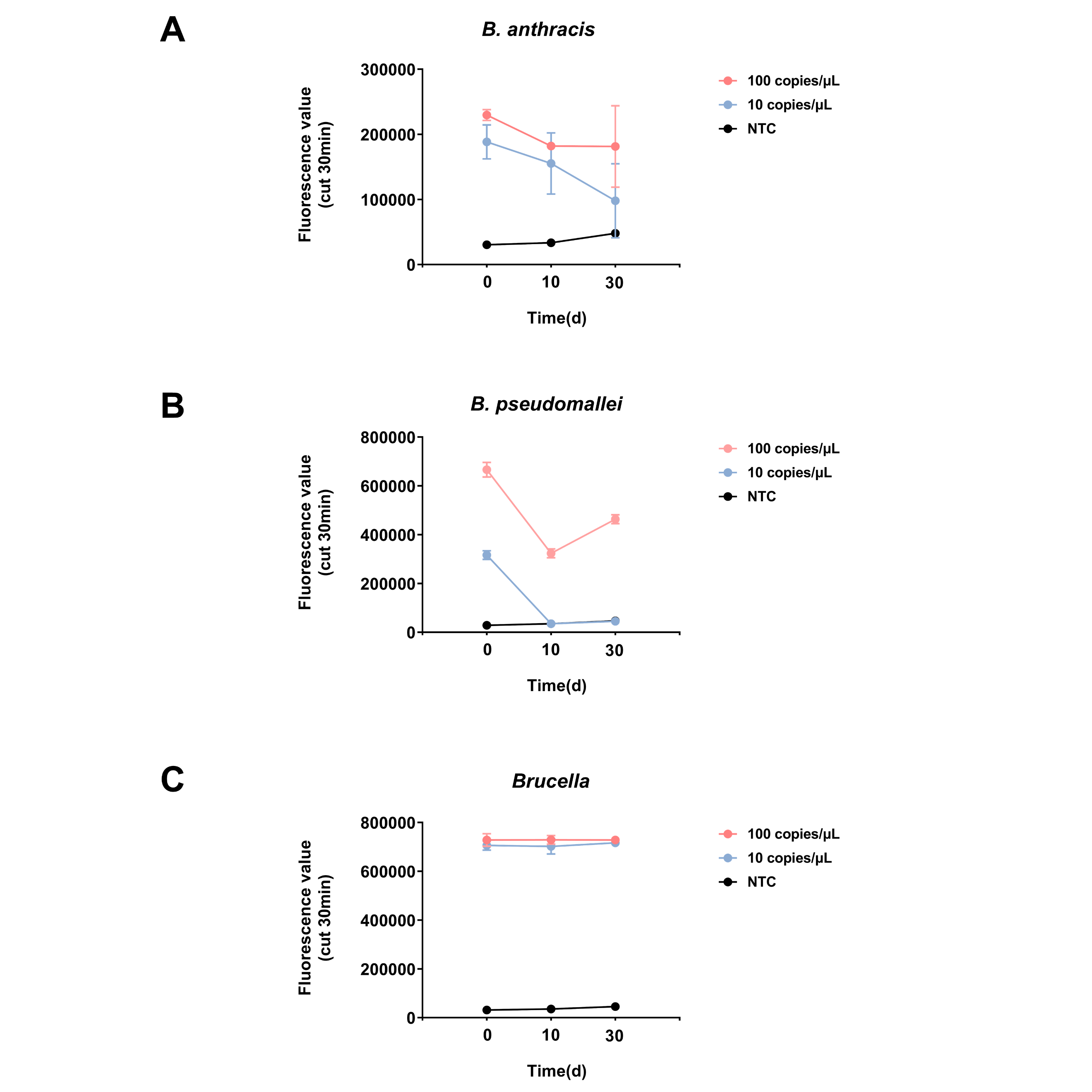

**Figure S4** Stability evaluation of the one‑step RPA/CRISPR lyophilized system, n=, 2mean ±SEM. (A) *B. anthracis*, (B) *B. pseudomallei* and (C) *Brucella*.

**Reference**

[1] J.H. Xu, X.R. Bai, X.L.L. Zhang, B. Yuan, L. Lin, Y. Guo, Y.J. Cui, J. Liu, H. Cui, X. Ren, J.L. Wang, Y. Yuan, Development and application of DETECTR-based rapid detection for pathogenic *Bacillus anthracis*, *Anal. Chim. Acta*. **2023**, 1247, 340891.

[2] J.X. Zhang, J.H. Xu, B. Yuan, X.D. Wang, X.H. Mao, J.L. Wang, X.L.L. Zhang, Y. Yuan, Detection of *Burkholderia pseudomallei* with CRISPR-Cas12a based on specific sequence tags, *Front. Public. Health*. **2023**, 11, 1153352.

[3] J.H. Xu, J.F. Ma, Y.W. Li, L. Kang, B. Yuan, S.Q. Li, J. Chao, L.H. Wang, J.L. Wang, S. Su, Y.Yuan, A general RPA-CRISPR/Cas12a sensing platform for *Brucella* spp. detection in blood and milk samples, *Sensors and Actuators: B. Chemical* **2022**, 364, 131864.
